# Sonic hedgehog signaling promotes basal epidermal *fibrillin 3* expression for zebrafish fin ray branching

**DOI:** 10.1101/2025.11.07.687116

**Authors:** Samuel G. Horst, Gabriel A. Yette, Astra L. Henner, Scott Stewart, Kryn Stankunas

**Affiliations:** Institute of Molecular Biology University of Oregon 297A Klamath Hall 1370 Franklin Blvd Eugene, OR 97403-1229; Department of Biology University of Oregon 297A Klamath Hall 1370 Franklin Blvd Eugene, OR 97403-1229

## Abstract

Zebrafish robustly regenerate amputated fins, including restoring branched bony ray skeletons. During fin outgrowth, *sonic hedgehog a* (*shha*)-expressing basal epidermal cells (bEps) signal to themselves and adjacent progenitor osteoblasts (pObs) to progressively split pObs into daughter ray pools. Collectively moving bEps pass through a *shha*-expressing state to define distal *shha*- positive bEp domains that themselves separate laterally ahead of ray branching. As such, Shh/Smo signaling may promote transient associations between bEps and pObs that gradually splits pOb pools. We used bulk RNA sequencing of caudal fin regenerates after Smoothened (Smo) inhibition followed by a CRISPant screen to identify *fibrillin 3* (*fbn3*) as a candidate Shh/Smo target gene for ray branching. Homozygous *fbn3* mutant zebrafish regenerated caudal fins with fewer branched rays and increased distances to branch points. Shh/Smo signaling was required specifically for full *fbn3* induction in bEps but not pObs. Double mutants showed that *fibrillin 2* (*fbn2*; *FBN2* ortholog) redundantly supports *fbn3*-promoted ray branching. Fbn1-containing microfibrils accumulated at the bEp/pOb interface in *fbn3* and, more so, *fbn3/2* mutant regenerating caudal fins. Shh/Smo signaling may activate bEp *fbn3* expression to locally modify microfibril abundance, facilitating bEp-pOb interactions and/or movements underlying ray branching morphogenesis.

## INTRODUCTION

Adult zebrafish fins robustly regenerate following amputation, including restoring their characteristic bony ray skeletons (reviewed in Sehring and Weidinger 2020). The well-studied caudal fin typically has 18 rays, with the 16 central-most rays discretely branching one or more times. Ray pattern, including branching, is re-established following caudal fin amputation by processes shared across median and paired fins. In all cases, discrete pools of progenitor osteoblasts (pObs) established after injury and residing within the distal blastema proliferate, organize, and differentiate to progressively regenerate bony rays.

Ray branch position is an emergent property arising from the gradual splitting of each ray’s pOb pool during fin outgrowth. Once pOb pools are divided, the branch point is reinforced by osteoblast-mediated “stitching” and osteoclast-driven resorption (Cardeira-da-Silva et al., 2022). Endocrine signals also influence branching position (Autumn et al., 2024; Harper et al., 2023). Sonic hedgehog (Shh) signaling specifically promotes pOb pool splitting during ray branching (Armstrong et al., 2017; Braunstein et al., 2021). *sonic hedgehog a* (*shha*) is expressed in distal domains of basal epidermal cells (bEps) adjacent to pObs at the leading edge of each regenerating ray (Armstrong et al., 2017; Hadzhiev et al., 2007; Laforest et al., 1998; Lee et al., 2009; Zhang et al., 2012). Shh signaling is active in both bEps and pObs throughout fin outgrowth – not only preceding overt ray branching (Armstrong et al., 2017; Braunstein et al., 2021). However, bEps collectively and continuously move distally (Braunstein et al., 2021). Therefore, the bEps comprising the *shha*-positive domains are constantly renewed, with individual bEps transiently expressing *shha* and responding with active Shh signaling only while passing alongside pObs. In contrast, pObs continuously respond to Shh signals until they re-differentiate and contribute to the extending rays. Collectively, *shha-*expressing bEp domains gradually split concurrently with the underlying pOb pools (Armstrong et al., 2017; Braunstein et al., 2021; Zhang et al., 2012). BMS- 833923 (BMS), a Smoothened (Smo) inhibitor that blocks Shh signaling and ray branching, prevents pOb pool but not *shha*-expressing bEp domain splitting (Armstrong et al., 2017; Braunstein et al., 2021). Further, BMS treatment prevents ray branching even after pOb splitting has initiated (Armstrong et al., 2017). Therefore, Shh/Smo-driven ray branching involves local Shh-driven transcriptional responses in both bEps and pObs that gradually splits pObs into daughter ray-forming pools.

pObs reside directly adjacent bEps, separated only by a distally-fragmented basement membrane through which the two cell types appear to make heterotypic contacts (Armstrong et al., 2017). This observation suggests that the continuous Shh/Smo signaling for ray branching could promote co-associations between pObs and moving bEps that gradually splits pObs into distinct pools. In further support, basement membrane disruption in *fraser extracellular matrix complex subunit 1* (*fras1*) mutants separates the bEp and pOb layers and disrupts ray branching even though Shh/Smo signal transduction remains intact (Robbins et al., 2023). Therefore, Shh signaling in pObs, bEps, or both populations may transcriptionally upregulate heterotypic cell adhesion proteins or extracellular matrix proteins that lead pObs to follow *shha*-expressing bEps comprising the splitting bEp domains. Identifying Shh/Smo targets would facilitate exploring these models or suggest alternative hypotheses how Shh/Smo promotes ray branching morphogenesis.

We used bulk RNA sequencing of acutely Shh/Smo-inhibited regenerating caudal fins to identify candidate Shh/Smo transcriptional targets. We targeted select genes expressed in bEps and/or pObs by a CRISPant screen to identify *thsd7ba* and *fbn3* (previously known as *fbn2b*; human *FBN3* ortholog) CRISPants with fin regeneration ray branching defects. Homozygous *fbn3* mutant fish showed delayed ray branching and lacked secondary branches during fin regeneration. Shh signaling remained active and *shha-*expressing bEp domains split normally in *fbn3* mutant fins, consistent with Fbn3 acting downstream of Shh/Smo signaling. *fbn3* expression was specifically reduced in *shha*-expressing bEps upon Smo inhibition. Double homozygous mutants for *fbn2* and *fbn3* had more severe ray branching defects. Surprisingly, Fbn1-containing microfibrils accumulated and/or formed altered conformations upon the loss of *fbn3* or *fbn3* with *fbn2*. We propose Shh signaling activates basal epidermal *fbn3* expression to locally suppress or modify microfibrils in a manner supporting pOb movements with dividing domains of Shh-expressing bEps. As such, Shh/Smo signaling helps produce a locally permissive environment that, together with additional transcriptional targets, promotes ray branching morphogenesis.

## RESULTS

### A CRISPR screen identifies candidate Sonic hedgehog signaling target genes promoting ray branching

The Shh/Smo signaling target genes in both Shh-producing basal epidermal cells (bEps) and adjacent pre-osteoblasts (pObs) that gradually drive pOb pool splitting and subsequent ray branching morphogenesis are unknown. We performed bulk RNA sequencing of 4-day post amputation (dpa) distal regenerates following an acute, 5-hour treatment with the specific Smo inhibitor BMS-833923 (BMS) to identify candidate transcriptional targets. BMS treatment downregulated 28 genes, including the universal Shh/Smo target genes *ptch1*, *ptch2*, *gli1*, and *hhip* (Fig. 1A-B; Supplemental Table 1). We cross-referenced these genes to our single cell RNA sequencing dataset (Lewis et al., 2023; Lewis et al., 2025) to identify genes expressed in the Shh/Smo-responsive basal epidermis and/or osteoblast clusters. 23 transcripts were represented between the two cell types, with 16 in basal epidermal cells and 15 in osteoblasts. The unbiased identification of positive controls and the short list of candidates enriched in the relevant cell types indicates the likely representation of key, direct target genes.

**Figure 1.**
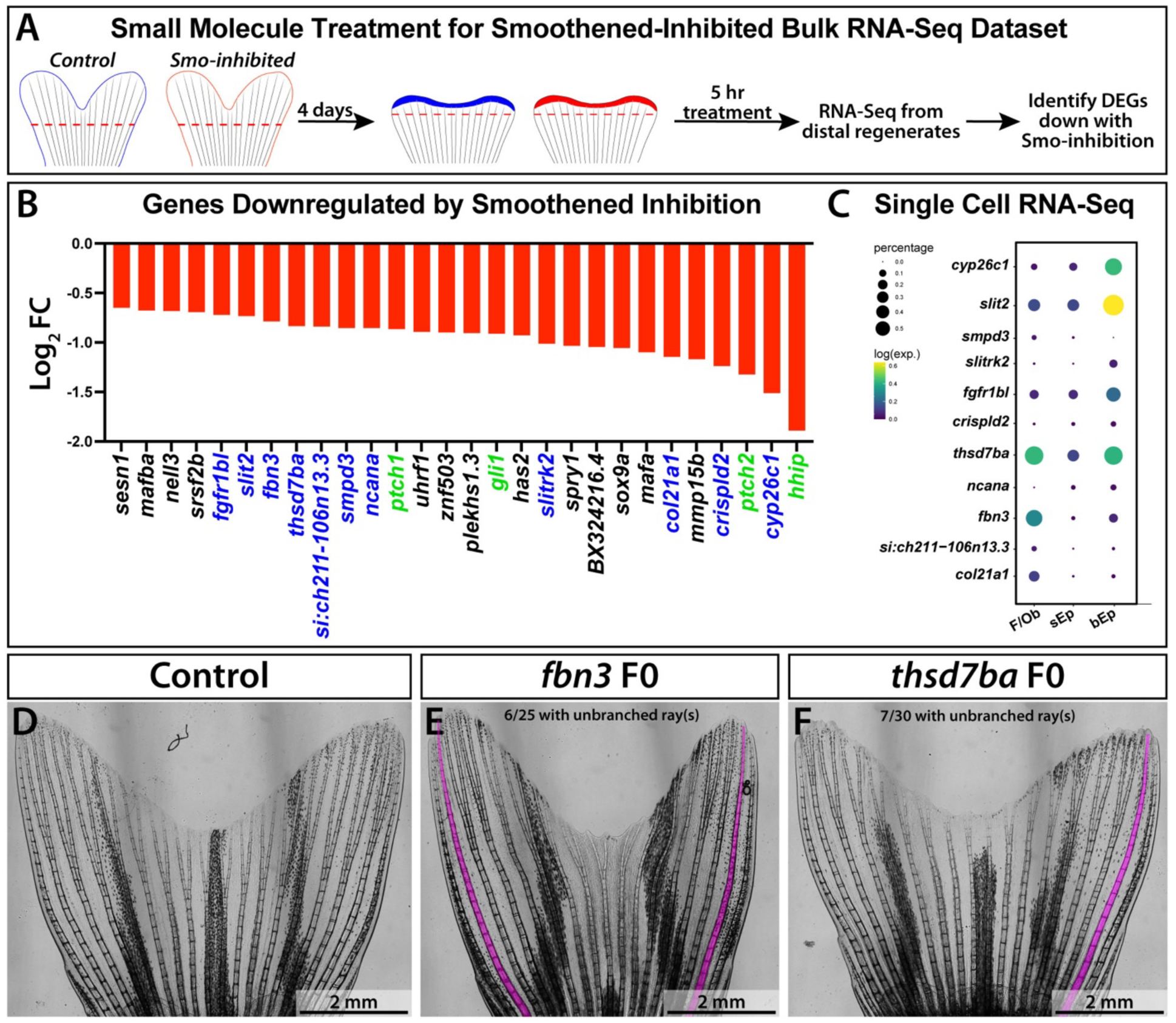
A CRISPant screen to identify genes downstream of Shh signaling-promoted ray branching. **(A)** Schematic of small molecule treatment and sample collection for bulk RNA-seq. Smoothened-inhibited fish were treated with BMS-833923 for the five hours immediately prior to tissue harvest. **(B)** Genes significantly downregulated in Smoothened-inhibited fin regenerates. Genes selected for the CRISPant screen are shown in blue; canonical Shh targets are shown in green. Log_2_ fold change (Log_2_FC) of transcript abundance is plotted on the y-axis. **(C)** Bubble plots showing expression enrichment of the genes selected for the CRISPant screen across fibroblast/osteoblast (F/Ob), superficial epidermis (sEp), and basal epidermis (bEp) clusters from a combined 3 and 7 dpa scRNA-Seq dataset (Lewis et al., 2023). **(D–F)** Whole mount caudal fin images of 27 dpa adult CRISPants for uninjected controls, *fbn3*, and *thsd7ba*. Rays with complete branching defects are highlighted in magenta.

Shh/Smo signaling might promote heterotypic cell associations between bEps and pObs that leads the pObs to follow splitting Shh-expressing basal epidermal domains (Armstrong et al., 2017; Braunstein et al., 2021). Therefore, we focused functional studies on the 11 candidate Shh/Smo target genes that encode transmembrane or extracellular matrix (ECM) proteins that could mediate such cellular co-movements (Fig. 1C). We performed a F0 CRISPant screen targeting each gene separately with two guide RNAs designed to disrupt early exons and generate null mutations. Adult zebrafish targeted for two genes, *thrombospondin domain containing protein 7ba* (*thsd7ba*) and *fibrillin 3* (*fbn3*), showed mosaic caudal fin branching defects. *fbn3* and *thsd7ba* CRISPant caudal fins were missing branchpoints in 6 of 25 fish and 7 of 30 fish, respectively (Fig. 1D-F). We established germline-transmitted mutant alleles *fbn3^b1468^ and thsd7ba^b1494^* and bred each to homozygosity. *fbn3* but not *thds7ba* homozygous mutant adults consistently showed delayed or absent ray branching in regenerated caudal fins (Fig. 1D-F). Therefore, we focused on *fbn3* as the best candidate Shh/Smo signaling target gene for ray branching morphogenesis.

### *fibrillin 3* promotes ray branching during caudal fin development and regeneration

*fbn3* was recently renamed from *fbn2b* as the ortholog of human *FBN3* based on phylogenetics and gene synteny (Supp. Fig. 1; Hu et al. 2023). *fbn3* is one of three zebrafish fibrillins (*fbn1*, *fbn2* [formerly *fbn2a*], and *fbn3*), each with a human paralog. The *fbn3^b1468^* allele (hereafter, *fbn3^-^*) has a 5 base pair deletion in exon 2 that produces an early stop codon. The highly truncated predicted product would have a disrupted N-terminal domain and exclude all downstream regions, including the EGF-like, TGF-ß binding, and integrin binding sites (Fig. 2A). *fbn3^-/-^* fish showed larval fin blistering phenotypes that match those of other *fbn3* mutant alleles *puff daddy* and *scotch tape* (Supp. Fig. 2, Gansner et al. 2008; Mellman et al. 2012). Nonetheless, *fbn3^-/-^*juveniles still developed caudal fins, albeit frequently with abnormal endochondral skeletons, including an indistinct hypural diastema and fused hypurals (Supp. Fig. 2). Most *fbn3^-/-^* fish survived to adulthood, developing to normal body lengths. The caudal fins of most *fbn3^-/-^* fish had significantly fewer caudal fin rays with correspondingly decreased average caudal fin widths (Fig. 2B-F). However, those rays developed and then regenerated to normal lengths (Fig. 2M). And, otherwise, *fbn3* mutant fish developed largely normal caudal fins with intact, segmented, and tapered rays, enabling loss-of-function studies of *fbn3* during fin ray branching.

**Figure 2.**
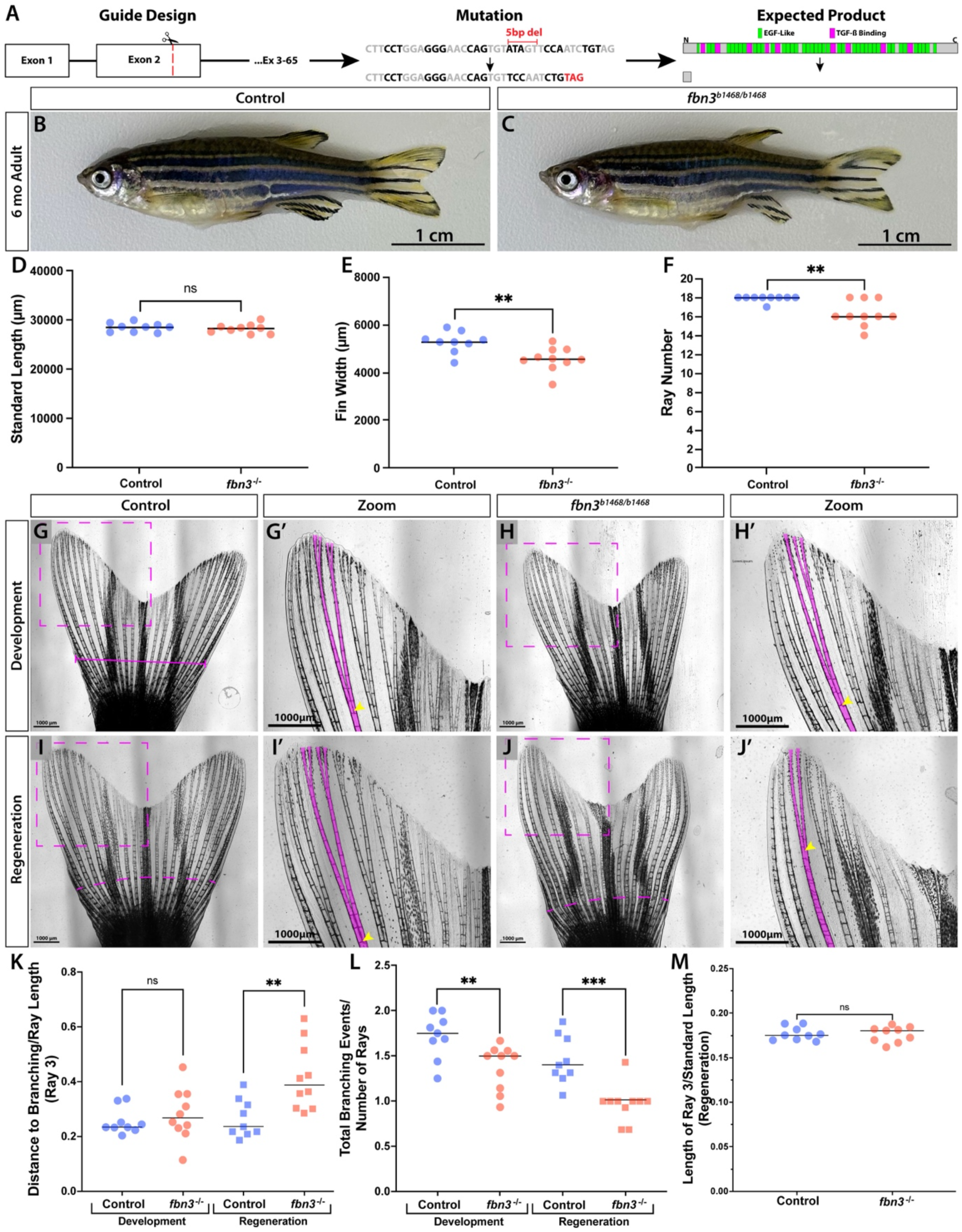
Fibrillin 3 supports ray branching during caudal fin regeneration. **(A)** Schematic showing the *fbn3* gRNA target site, resulting germline mutation with premature stop codon, and predicted truncated protein. **(B, C)** Whole mount images of sibling control and *fbn3^-/-^* adult zebrafish. **(D-F)** Dot plot graphs comparing standard length, caudal fin width (intersecting the longest procurrent rays), and caudal fin ray number of control and *fbn3^-/-^*fish. **(G-H’)** Pre- amputation caudal fin images from sibling control and *fbn3^-/-^* animals. The magenta line in (G) indicates fin width measurement. In (G′, H′), ray 3 is outlined in magenta; yellow arrowheads mark branchpoints. **(I-J’)** Regenerated caudal fin images of the same animals shown in (G-H’) imaged 30 days post amputation. The magenta dashed lines denote amputation planes. **(K-M)** Dot plot graphs comparing ray branching and outgrowth in regenerated caudal fins of control and *fbn3^-/-^* fish. (K) The ratio of branchpoint distance to total ray length during development and regeneration, measured from the longest procurrent rays (pre-amputation) or from the amputation plane (post- regeneration). (L) The number of branching events per branching ray during development and regeneration (principal peripheral rays excluded). (M) Ratio of ray 3 length to standard length. Data from three independent experiments (n = 9 controls, n = 10 mutants). All data points and means are shown. Significance determined by one-tailed, unpaired Student’s *t*-test: *P < 0.05 (*), P < 0.005 (**), P < 0.0005 (***)*.

Caudal fins of *fbn3^-/-^* adult fish had fewer branching events per ray albeit with an insignificant change in the average distance to a representative ray’s (ray 3) first branchpoint (Fig. 2G-H’, K, L). Branching abnormalities were markedly more severe in *fbn3^-/-^* regenerated fins, showing both fewer branching events per ray and a significant increased distance to ray 3’s first branchpoint (Fig. 2I-L). We conclude that Fbn3, while not essential, promotes ray branching during fin outgrowth with a more pronounced role during regeneration.

We used the *Tg(−2.4shha:gfpABC)^sb15^* (*shha:GFP*) reporter to find *shha*-expressing bEp domains of regenerating *fbn3^-/-^* caudal fins still split over a 3 to 6 dpa time course (Fig. 3A-F). We used *TgBAC(ptch2:kaede)^a4596^* (*ptch2:Kaede*) fish to monitor Shh output in both *shha*-expressing bEps and distal pObs of *fbn3* mutants (Armstrong et al., 2017; Braunstein et al., 2021; Huang et al., 2012; Laforest et al., 1998; Quint et al., 2002). We photoconverted Kaede-expressing cells of caudal fin distal regenerates of *fbn3^-/-^* and control fish at 3 dpa and then re-imaged at 20 hours post-conversion (hpc). Distal bEps and pObs produced new, unconverted Kaede in both *fbn3^-/-^* mutants and control siblings at 20 hpc (Fig. 3G-K’). These results collectively show that Fbn3 does not modulate *shha* expression, *shha*-expressing bEp domain splitting, or Shh/Smo transcriptional output, consistent with Fbn3 supporting ray branching as a downstream Shh/Smo target gene.

**Figure 3.**
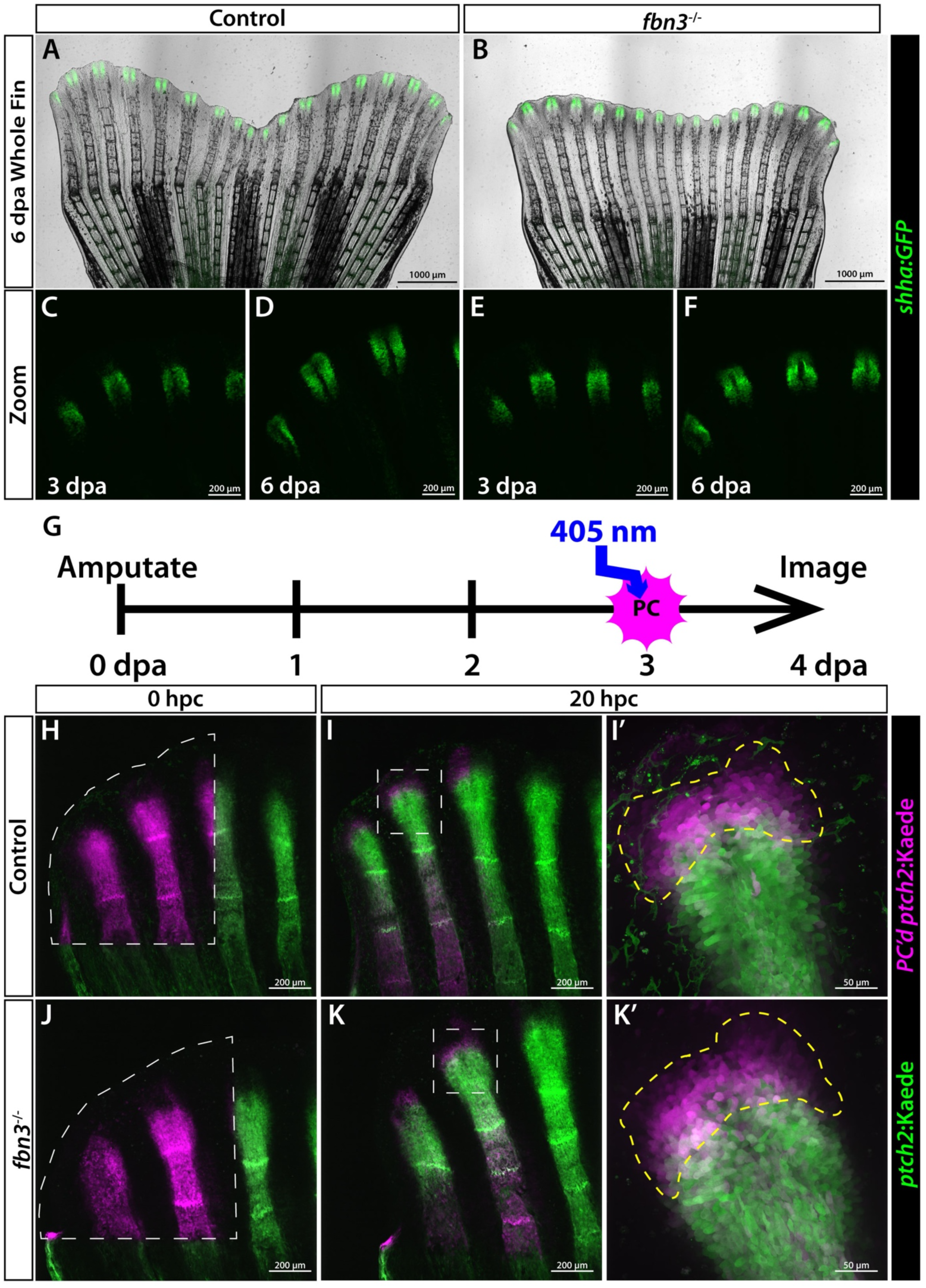
Fibrillin 3 does not mediate Sonic hedgehog signaling or splitting of *shha*- expressing bEp domains during fin regeneration. **(A, B)** Wholemount images of 6 dpa regenerated caudal fins from sibling control and *fbn3^-/-^* fish carrying the *shha:GFP* transgene. **(C- F)** Wholemount images showing *shha:GFP* domains at regenerating dorsal caudal fin rays of (C, D) control and (E, F) *fbn3^-/-^* fish at 3 and 6 dpa (n=9 mutants and n=5 controls). **(G)** Schematic of *ptch2:kaede* photoconversion experiment. Distal regions of dorsal rays are photoconverted at 3 dpa and imaged immediately and again at 4 dpa. **(H-K)** Overlaid confocal fluorescent images of (H, I) control and (J, K) *fbn3^-/-^* regenerating caudal fins at 0 hours post conversion (hpc) (3 dpa; dashed lines indicate photoconverted regions; H, J) and then 20 hpc (4 dpa; I, K). (I’, K’) Zoomed views of ray 2 at 20 hpc. Unconverted Kaede is green and photoconverted Kaede is magenta. Photoconverted bEps that have migrated past the active Shh signaling zone are outlined in yellow (n=5 mutants and n=5 controls).

### Shh signaling promotes *fbn3* expression specifically in *shha*-expressing bEps

Shh signaling is active in both *shha*-expressing bEp and distal pObs (Armstrong et al., 2017; Braunstein et al., 2021). Therefore, Shh signaling could promote *fbn3* expression in bEps, pObs, or both populations. We used RNAScope hybridization to monitor *fbn3* transcripts in 4 dpa *shha:GFP*-expressing longitudinal fin sections under normal conditions and after a 4-hour acute BMS treatment (Fig. 4A-B’). Control sections revealed that *fbn3* is normally expressed in distal, *shha*-expressing bEps, pObs, and both proximal and distal fibroblasts. The brief Smo inhibition specifically decreased *fbn3* levels in *shha*-expressing bEps (Fig. 4C). Therefore, Shh signaling activates *fbn3* transcription in bEps but not the adjacent Shh/Smo-responsive pObs or underlying blastema mesenchyme.

**Figure 4.**
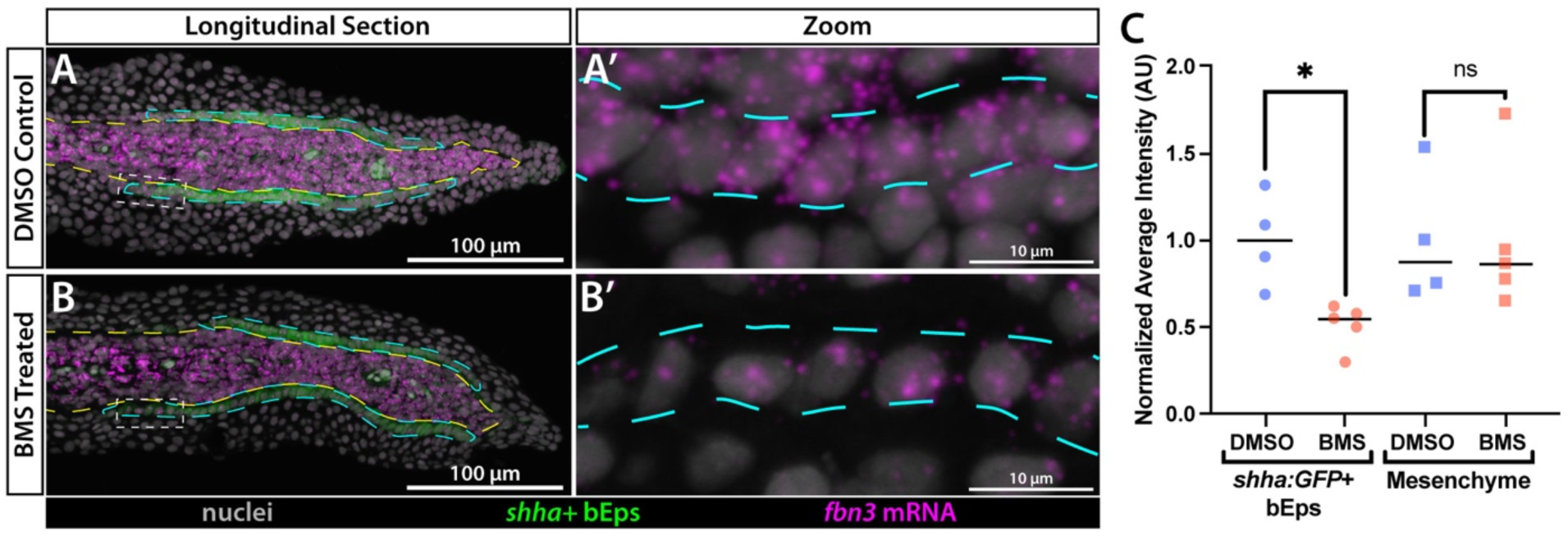
Shh signaling promotes *fbn3* expression specifically in *shha*-expressing bEps of regenerating fins. **(A-B)** Confocal fluorescent images of longitudinal 4 dpa regenerating caudal fin sections from *(A)* DMSO and (B) BMS-833923 treated fish. RNAScope-stained *fbn3* transcripts are shown in magenta, anti-GFP antibody stained *shha:GFP*-positive basal epidermal cells (bEps) in green, and Hoechst-stained nuclei in grayscale. The fibroblast and osteoblast populations are outlined with a yellow dashed line, and the *shha:GFP* domain is highlighted in cyan. White dashed boxes in (A, B) indicates the zoomed regions in (A′, B’). **(C)** Dot plot graphs showing normalized *fbn3* transcript intensity across *shha:GFP*-expressing bEp domains and mesenchyme (outlined in A). Each point represents a section from one fish (n = 4 control, n = 4 mutant). Significance determined by two-tailed unpaired Student’s *t*-test with a Bonferroni correction; *P < 0.025 (*)*.

### Fibrillin 3 suppresses microfibril formation in the distal blastema

Mutants for the *fras1* Fraser Complex gene also show reduced ray branching associated with disrupted interconnectivity between the basal epidermal and underlying mesenchymal layers (Robbins et al., 2023). In contrast, anti-Laminin staining of 4 dpa *fbn3^-/-^*regenerating caudal fin sections showed an intact basement membrane associated with both overlying bEps and underlying pObs (Supp. Fig.4). Further, alpha-catenin staining indicated the basal epidermis maintained adherens junctions and epithelial characteristics in the absence of *fbn3* (Supp. Fig. 4). Therefore, ray branching defects in *fbn3* mutants likely reflects a local extracellular matrix organization role for Fbn3 in ray branching morphogenesis.

We used the well-established JB3 anti-Fibrillin antibody to monitor microfibril distribution in regenerating caudal fins of wildtype and *fbn3^b1468/b1468^* fish (Wunsch et al. 1994). Microfibrils were prominent proximally both between hemi-rays and within inter-ray spaces of control 4 dpa regenerating fin sections (Fig. 5C). In contrast, microfibrils were largely absent in distal regions, including at the bEp and pOb interface (Fig. 5A, D. In *fbn3^-/-^* mutants, proximal fibrillin staining was stronger but more punctate, with fewer extended microfibrils (Fig. 5C, E). Further, we observed abundant microfibrils distally between *shha*-expressing bEps and pObs uniquely in *fbn3^-/-^* animals (Fig. 5B, F). Transverse sections highlighted that the ectopic microfibrils extended perpendicularly from the bEp/pOb interface and into the blastema mesenchyme (Fig. 5F-F’’). We conclude the JB3 antibody is likely recognizing Fbn1and/or Fbn2-containing microfibrils given the elevated signal upon *fbn3* loss. Further, Fbn3 expression modifies the levels and size of microfibrils in both proximal, differentiated fin tissue and in the distal regenerate where branching morphogenesis actively occurs.

**Figure 5.**
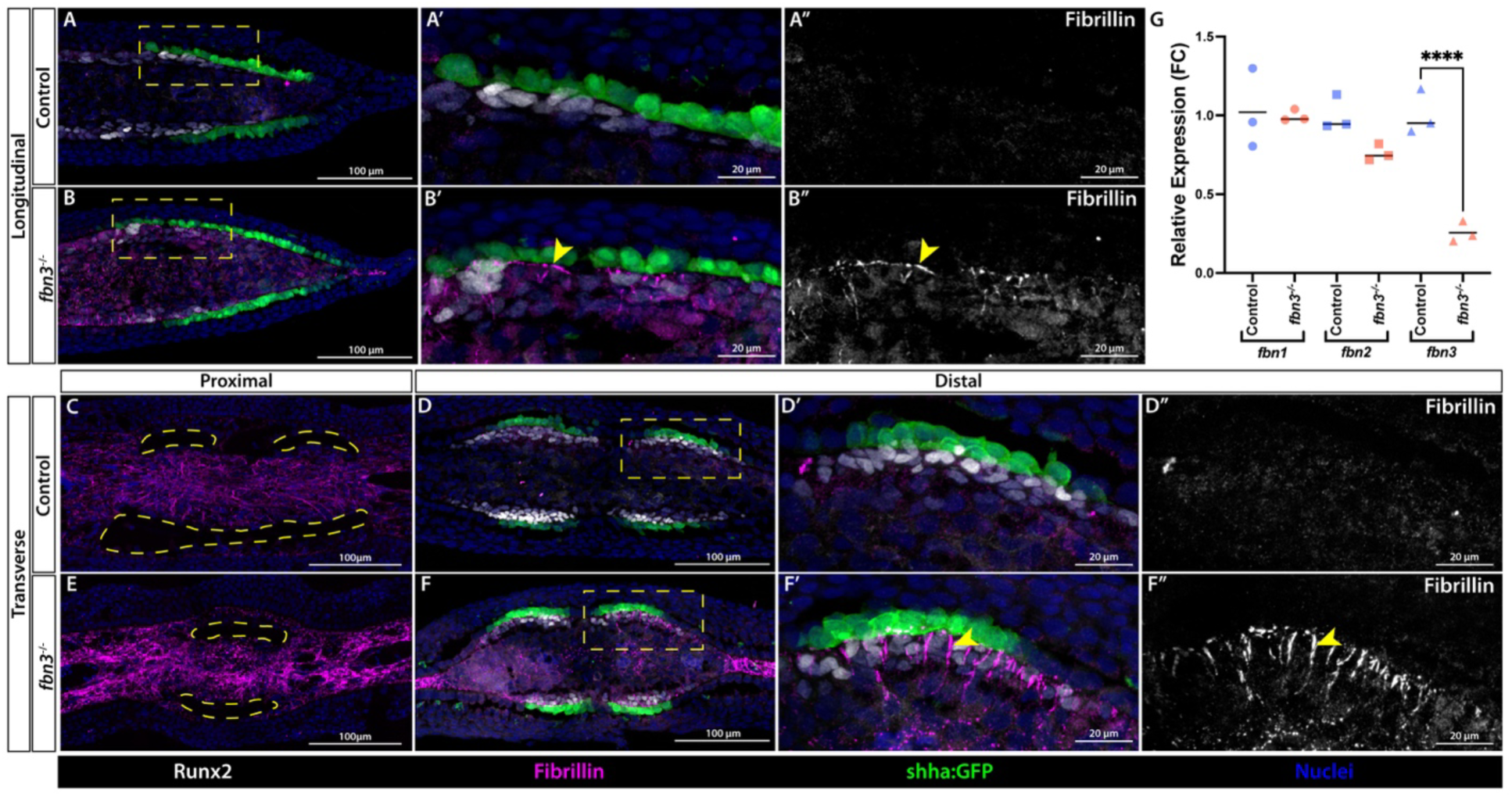
Fibrillin 3 restrains microfibril production in regenerating fins, including at the distal regenerate pOb/bEp interface. **(A-F)** Confocal immunofluorescence images of sections from 4 dpa caudal fin regenerates. Both control and clutchmate *fbn3^-/-^*fish carry the *shha:GFP* transgene. For overlay images, Fibrillin is in magenta, *shha:GFP*-expressing basal epidermal cells in green, Runx2-expressing pObs in grayscale, and Hoechst-stained nuclei in blue. The yellow dashed boxes in (A, B, D, F) mark the zoomed regions shown in overlay (A′, B’, D’, F’) or with Fibrillin alone in grayscale (A″, B”, D”, F”). (A, B) Longitudinal and (C-F) transverse sections are shown. The yellow arrowheads in (B′, B’’, F’, F”) highlight prominent microfibrils found distally only in *fbn3^-/-^* regenerates. The yellow dashed outline in (C) marks forming bone in the proximal regenerate section. Longitudinal section staining was repeated three times, each time using three mutant and three control sections. Transverse section staining was repeated twice using three mutant and three control sections. **(G)** Dot plot graph showing the relative expression of *fbn1*, *fbn2*, and *fbn3* by RT–qPCR using cDNA from three pools of 4 dpa caudal fin regenerates (*fbn3^-/-^* and age-matched controls). Expression was normalized to *rpl8*; statistical significance was assessed by one-way ANOVA followed by Bonferroni’s multiple-comparison test: *P < 0.00005 (****)*.

We used RT-qPCR of 4 dpa caudal fin regenerates to investigate if the apparent increase in microfibrils in *fbn3* mutants resulted from upregulated *fbn1* and/or *fbn2* transcripts. However, *fbn1* and *fbn2* expression were unchanged while *fbn3* transcripts were decreased, presumably due to nonsense-mediated decay (Fig. 5G). Therefore, the increased microfibril staining in *fbn3* mutants likely does not reflect genetic compensation by *fbn1* or *fbn2*. Rather, this result further supports that Fbn3 either suppresses formation of Fbn1 and or Fbn2-containing microfibrils or alters microfibril structure or composition in a manner enhancing antigenicity.

Shh/Smo signaling upregulation of basal epidermal *fbn3* could suppress or modify microfibril formation in distal regenerating tissue as part of ray branching morphogenesis. To test this, we stained 48-hour BMS-treated 4 dpa regenerating caudal fin sections with the JB3 anti-Fibrillin antibody. BMS-treated samples showed increased Fibrillin staining at the bEps and pObs interface compared to DMSO controls (Supp. Fig. 5). However, the effect was more subtle than with *fbn3* loss-of-function and rarely included longer ectopic microfibrils, consistent with Shh/Smo activating *fbn3* only in bEps and not blastemal cells. The stronger microfibril phenotype in *fbn3* mutants suggests that *fbn3* expression from both Shh-responsive and -insensitive sources contributes to microfibril modulation. Conversely, the more severe branching defects following BMS treatment indicate that Shh/Smo signaling promotes ray branching through additional targets beyond *fbn3*.

### Fibrillin-2 and Fibrillin-3 contribute to ray branching morphogenesis

Fbn2 and Fbn3 co-expression in distal fin regenerates, their high amino acid similarity, and shared functionality in other contexts suggested they could act redundantly, explaining delayed but not abolished branching in *fbn3* mutants. We generated a *fbn2^b1481^* (*fbn2^-^*) allele with a 5 bp deletion and therefore frameshift and early stop codon in exon 2. *fbn2^-/-^* fish did not have notable fin development or regeneration defects, including ray branching pattern changes (Supp. Fig. 6). *fbn3^-/-^*; *fbn2^-/-^* fish were viable and fertile, and largely phenocopied *fbn3^-/-^* single mutants, including developing caudal fins with subtle ray branching defects. However, *fbn3^-/-^*; *fbn2^-/-^* fish regenerated caudal fins with even further delayed ray branching and fewer branching events per ray (Fig. 6A- D). As such, *fbn2* and *fbn3* redundantly promote ray branching but with the major functional contribution from Fbn3.

**Figure 6.**
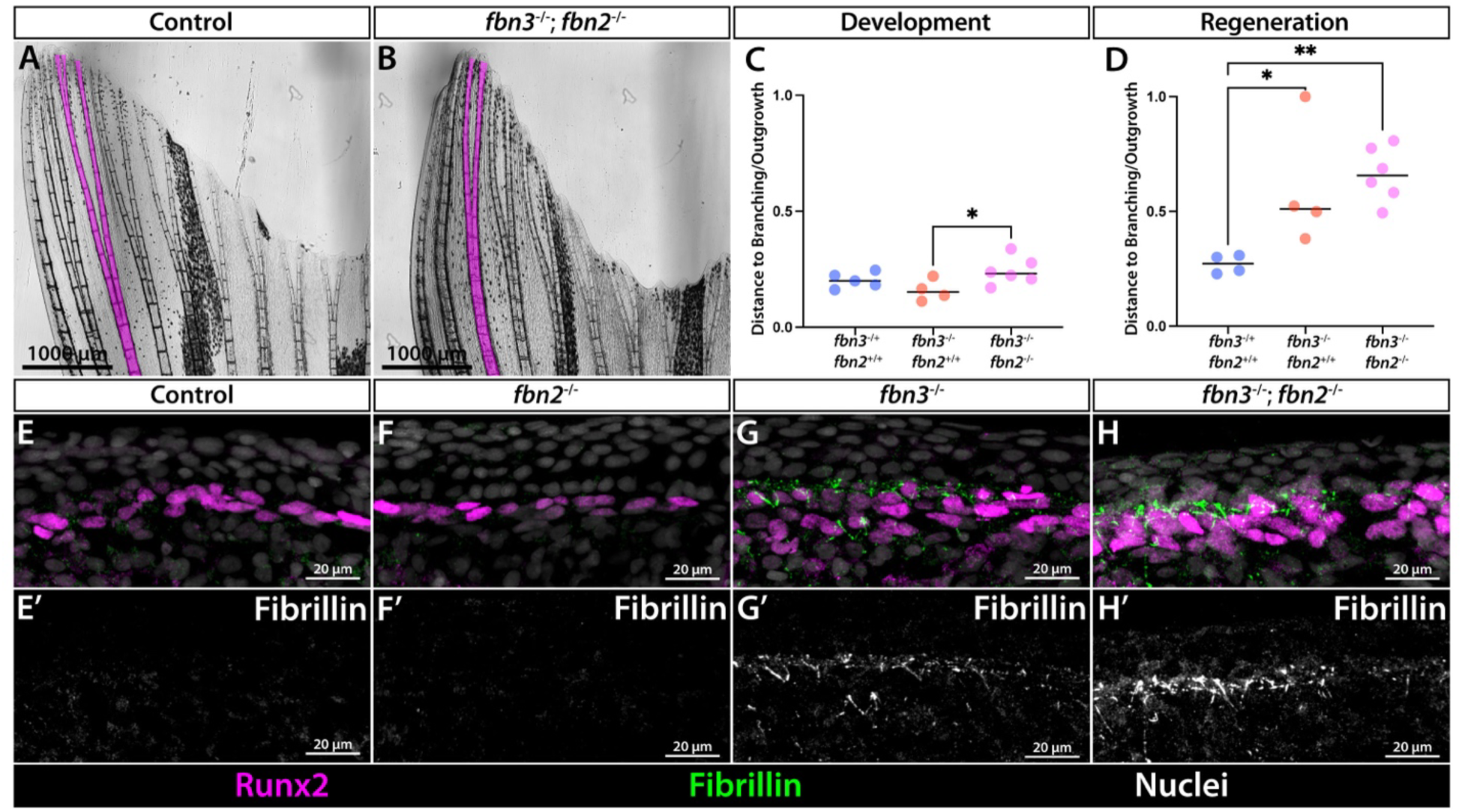
Fibrillin 2 supports Fibrillin 3-promoted ray branching and microfibril organization in regenerating fins. **(A-B)** Images of 30 dpa distal caudal fin regions from a control and *fbn3^-/-^*; *fbn2^-/-^* clutchmate fish. Ray 3 is highlighted in magenta. **(C, D)** Dot plot graphs showing the ratio of branchpoint distance to total ray outgrowth during development and regeneration. All data points and means are shown. Significance determined by one-way ANOVA with Tukey’s post-hoc test: *P* < 0.05 (*), *P* < 0.005 (**). Regeneration and development experiments were repeated with independent animals. (A-D) are also shown with extended data in Supplemental Figure 3. **(E-H)** Confocal immunofluorescent images of longitudinal 4 dpa distal regenerate sections from control, *fbn2*^-/-^, *fbn3^-/-^*, and *fbn3^-/-^*; *fbn2^-/-^* fish. Fibrillin (presumably, Fibrillin 1) is in green, Runx2-marking pObs is in magenta, and Hoechst-stained nuclei are in grayscale. (E’-H’) Corresponding grayscale images showing Fibrillin staining alone. (E-H) are also shown with extended data in Supplemental Figure 7. Fibrillin staining on matched controls and *fbn3^-/-^*; *fbn2^-/-^* samples was repeated twice, each on sets of 3 animals per group.

We stained *fbn3^-/-^*; *fbn2^-/-^* and control 4 dpa caudal fin sections with JB3 to determine if both Fbn2 and Fbn3 contribute to microfibril modulation. The microfibril staining further increased in distal regenerate tissue of double mutants (Fig. 6E-H’, Supp. Fig. 7). Therefore, Fbn1- containing microfibrils either accumulate, enlarge, and/or become antigen-exposed in the absence of combined Fbn3 and Fbn2. We conclude Fbn3/Fbn2 restricts and/or alters microfibril formation in regenerating fins. Shh/Smo signaling contributes by inducing *fbn3* in distal basal epidermal cells. The Fbn3-altered extracellular matrix enables bEp and/or pOb movements that lead pObs into split pools for ray branching morphogenesis.

## DISCUSSION

### Fbn3 establishes an ECM environment supporting ray branching

We identify *fbn3* as a downstream Shh/Smo target in basal epidermal cells and a positive regulator of ray branching in regenerating zebrafish fins. Ray branching centrally reflects the progressive splitting of pOb pools, a process influenced by dynamic interactions with migrating bEps. Shh/Smo signaling promotes pOb movements and pool separation, potentially through transient heterotypic contacts with *shha*-expressing bEps (Armstrong et al., 2017; Braunstein et al., 2021). Fbn3, induced by Shh/Smo in bEps but also expressed in blastema cells, likely supports ray branching by influencing the presence and/or composition of microfibrils at the interface between *shha*-expressing bEps and pObs.

Fibrillins are large extracellular glycoproteins with EGF-like, TGF-β binding, integrin binding, and unique N- and C-terminal domains (Jensen et al., 2012). They form homo- or heterotypic extracellular microfibrils through N- to C-terminal linkages between monomers (Lin et al., 2002; Charbonneau et al., 2003; Kielty et al., 2005). In wildtype regenerating fins, microfibrils are largely absent from the distal blastema. In contrast, *fbn3* mutants show increased and structurally altered microfibril staining in the distal outgrowth zone. These are likely Fbn1- containing microfibrils given double *fbn3*; *fbn2* null mutants show even stronger microfibril accumulation. Further, the changes are not accompanied by increased *fbn1* or *fbn2* transcript levels, suggesting that Fbn3 modulates Fbn1-containing microfibril formation or structure rather than gene expression. The simplest explanation is that Fbn3 (with Fbn2) inhibits the formation of Fbn1 microfibrils. The lower overall distal Fbn1 levels in wildtype vs. *fbn3^-/-^* mutants could then result from degradation of unincorporated Fbn1 molecules by matrix metalloproteinases (Ashworth et al., 1999).

*fbn3* is expressed in both proximal and distal regenerating fin tissues. While microfibrils increase in both regions in *fbn3* mutants, the accumulation is more pronounced and structurally distinct in the distal regenerate. Proximal microfibrils, located both within and in between maturing bony rays, may provide structural stability in differentiated tissue. Similar, stabilizing Fbn1 microfibrils could then form in the absence of Fbn3 within the space between distal bEps and pObs. pObs might adhere to these ectopic distal Fbn1 microfibrils, restraining their movements with splitting *shha*-expressing bEp domains. Under this model, elevated distal levels of Fbn3 (with Fbn2) normally suppress Fbn1 microfibril deposition to produce an environment permissive to lateral pOb migration and ray branching (Supp. Fig. 8). A similar concept was recently proposed for tail bud development, where Fbn3 promotes ECM “fluidity” favoring progenitor cells (Genuth et al., 2025). Consistent with the possibility of Fbn cross-antagonism, recombinant Fbn1 and Fbn2 fragments bind at high affinity in a head-to-tail manner (Lin et al., 2002) and microfibrils can contain both Fbn1 and Fbn2 (Lin et al., 2002; Charbonneau et al., 2003). Further, co-culture of fibroblasts with cells expressing ectopic N-terminal Fbn2 reduces Fbn1 microfibril deposition (Charbonneau et al., 2003). Together, these findings suggest Fbn2/3 antagonism of Fbn1 could represent a broader mechanism for ECM modulation in other developmental, disease, and regenerative contexts.

An alternative explanation emerges if the elevated JB3 antibody signal in *fbn3* and *fbn3/fbn2* mutants reflects an altered microfibril structure that enhances antigen accessibility rather than an increase in microfibril abundance. In support, Fbn1 can mask Fbn2 epitopes within microfibrils of mouse tissue (Charbonneau et al., 2010). Therefore, Fbn3 may similarly influence the antigenicity of Fbn1-containing microfibrils. Under this scenario, specialized Fbn3-containing microfibrils may direct pOb movements for ray branching in addition to or distinct from a broader Fbn3 role in suppressing Fbn1 microfibrils. Consistent with this notion, Fibrillins interact with many proteins, including latent TGF Binding Proteins, microfibril-associated glycoproteins, Elastins, and Integrins that could produce a local ECM conducive to morphogenetic cell interactions (Bax et al., 2003; Jensen et al., 2001; Kielty et al., 2005; Thomson et al., 2019; Zilberberg et al., 2012). Notably, microfibrils promote cell adhesion and migration through their integrin binding sites (Bax et al., 2003; Chaudhry et al., 2007; Godwin et al., 2023; Nistala et al., 2010). Specific antibodies recognizing each zebrafish Fibrillin as well as examining pOb movement dynamics in *fbn2/fbn3* mutants would enable distinguishing Fibrillin roles, Shh-regulated or otherwise, in ray branching morphogenesis.

### Shh signaling activates distinct transcriptional programs in bEps and pObs

Shh signaling activates *fbn3* expression in the *shha*-expressing bEps but not in adjacent pObs. In contrast, Shh signaling activates other targets like *ptch2* in both cell types. Therefore, Shh/Smo signaling coordinates distinct molecular programs to support branching morphogenesis. By extension, *fbn3* clearly is not the sole target of Shh signaling effecting ray branching. First, Smo inhibition only modestly and locally reduced *fbn3* transcripts in bEps once assessed by in situ hybridization. Indeed, the major contribution of Fbn3 (and Fbn2) for ray branching may be Shh/Smo-independent and even outside of bEps. Second, *fbn3* and *fbn2* double mutant branching defects are less severe than those caused by Smo inhibition, even though microfibril accumulation is more dramatic. Therefore, additional Shh target genes, especially in pObs, likely promote adhesion or associations between pObs and bEps or modulate local environments that guide branching morphogenesis movements. The 22 additional differential expressed genes expressed in pObs and/or bEps remain candidates. This includes *thsd7ba*, which showed branching defects when targeted in CRISPants but not once germline-targeted, possibly reflecting genetic compensation. Identifying additional target genes may require higher resolution transcriptomics to isolate Shh/Smo-dependent transcripts in the small group of cells within regenerating fins with active Shh signaling.

### Fibrillin paralogs in zebrafish and implications for skeletal patterning

Disruption of fibrillin proteins causes human fibrillinopathies marked by skeletal abnormalities, suggesting conserved mechanisms with those underlying impaired ray branching in zebrafish fins. For example, mutations in *FBN1* cause Marfan syndrome, while *FBN2* mutations underlie congenital contractural arachnodactyly. *FBN3* is less well-characterized, likely due to its conspicuous absence from the mouse genome (Supp. Fig. 1; Corson et al. 2004). Zebrafish retains all three fibrillin paralogs, providing a unique system to parse paralog roles. Interestingly, zebrafish *fbn3* shares synteny with *FBN3* yet the protein has higher amino acid sequence similarity with FBN2 (Supp. Figure 1, Bateman et al. 2025; Harrison et al. 2024). Further, we show Fbn2 and Fbn3 have at least partially redundant roles in modulating microfibril production to support ray branching. Across species and contexts, Fbn2 and Fbn3 may function near interchangeably, but with variable roles or importance dependent on evolved spatiotemporal expression patterns. Relatively high *fbn3* expression could explain the predominant *fbn3* and partially redundant *fbn2* roles during zebrafish fin skeletal development and regeneration. Fbn1, which accumulates in *fbn3* mutants to seemingly disrupt cell behaviors of branching, also is normally expressed proximally where it may serve additional roles in zebrafish fin and skeletal regeneration. Regardless, zebrafish offers a promising system to explore distinct, shared, and interacting roles of fibrillin molecules to produce varied microfibril environments affecting skeletal patterning and organization.

## MATERIALS AND METHODS

### Zebrafish

*Danio rerio* zebrafish were maintained in 28–29 °C flowing water in the University of Oregon Aquatic Animal Care Services (UO AqACS) fish facility. Details of fish housing and feeding from larval stage through adulthood were detailed previously (Braunstein et al., 2021). Staff carried out feedings twice a day. Housing densities for fish of each developmental stage are also described in Braunstein et al. 2021. The following lines were used: wildtype AB, *TgBAC(ptch2:Kaede)^a4596^* (Huang et al., 2012), *Tg(−2.4shha:gfp:ABC)^sb15^* [previously known as *Tg(−2.*2shh*:gfp:ABC)*] (Ertzer et al., 2007; Shkumatava et al., 2004), *thsd7ba^b1494^, fbn3^b1468^*, and *fbn2^b1481^* . The University of Oregon Institutional Animal Care and Use Committee (IACUC) approved zebrafish experiments.

### BMS-833923 treatments

BMS-833923 (Cayman, 16240) was dissolved in DMSO (25 mg/mL), aliquoted, and stored at – 80 °C until use. For treatments, aliquots were diluted to 2.5–5 µM in 500 mL facility water; controls received equal volumes of DMSO. To account for batch variability, *ptch2:Kaede* fish were treated with new BMS stocks, photoconverted, and reimaged to calibrate dosing as described previously (Braunstein et al., 2021). Paraffin sections for *fbn3* in situ analysis were collected after a 4 h acute treatment at 4 dpa. Frozen sections were collected after a 24 h treatment (2–3 dpa) followed by a 4 h pulse at 4 dpa.

### RNA-Sequencing

*shha:GFP* heterozygous fish were injected intraperitoneally at 4 dpa with either 40 mg/kg BMS- 833923 (Cayman, #16240) in injection buffer (50% PEG-400, 5% Propylene Glycol, 0.5% Tween-80) or an equal volume of injection buffer. Five hours later, distal regenerates including and beyond the shha:GFP domains were collected in pools of four (three sets each for control and treated groups). RNA was extracted using Trizol, phase-separated with chloroform, precipitated with glycogen/isopropanol, washed in 70% ethanol, and resuspended in nuclease-free water. Libraries were prepared with the KAPA mRNA Seq kit and sequenced (Illumina HiSeq 4000, single-end). Reads were aligned to the zebrafish genome (GRCz11) with Tophat (Trapnell et al. 2009), quantified with HTSeq (Anders et al. 2015), and differential expression was assessed with EdgeR (Chen et al. 2025).

### CRISPR mutagenesis

CRISPR guides were designed with CHOPCHOP (Labun et al., 2019) to target early exons and induce frameshift mutations. Single-stranded oligos were ordered, amplified with a generic structural oligo, and transcribed in vitro using the MEGAscript T7 kit (Thermo Fisher, AM1333). Guide RNAs were purified (Zymo Clean & Concentrator-5, R1013) and prepared at 800 ng/µL with Phenol Red, nuclease-free water, and 100 ng/µL TrueCut HiFi Cas9 (Thermo Fisher, A50576) for injection into AB embryos at the one-cell stage. Embryos were raised under standard conditions. The *fbn3^b1468^*allele (5 bp deletion in exon 2) and *fbn2^b1481^* allele (5 bp deletion in exon 2) introduce early stop codons and generate null mutations. *fbn3^b1468^* heterozygotes displayed no phenotype, so heterozygotes and wildtype siblings were pooled as controls in subsequent experiments.

### Live microscopy

Adult Zebrafish were imaged in a walled coverslip (Cellvis Chambered Coverglass System #1.5 High Performance Cover Glass) after anesthesia using 168 mg/L Tricaine. Images were acquired on a Nikon Eclipse T*i-*E with Yokogawa CSU-W1 spinning disk attachments. Images were processed in Fiji and Adobe Photoshop with identical adjustments applied to all images in a given experiment. Live fish were euthanized or returned to the flowing water system immediately after imaging.

### Ray and fin measurements

In unamputated fins, ray length to the first branch point was measured from a reference line connecting the tips of the longest dorsal and ventral procurrent rays. This line corresponds to the amputation plane. In regenerated fins, ray length and distance to the first branch point were measured from the same reference line to the distal tip and first branched segment, respectively. Standard length was measured from head to the base of the caudal fin (Parichy et al., 2009).

### Alcian Blue/Alizarin Red Staining

Fish were sacrificed at 22 days post fertilization in Tricaine, then fixed in 4% PFA/PBS for an hour. Next, they were Rinsed in 50% EtOH before staining overnight in a solution of 0.04% Alcian Blue, 0.01% Alizarin Red, and 10 mM MgCl_2_ in 80% EtOH. Fish were then washed in 10mM MgCl_2_ in 80% EtOH, rehydrated by ethanol series, then bleached in 3% H_2_O_2_/0.5% KOH until Alizarin cleared sufficiently. Finally, fish were rinsed and stored in 50% Glycerol + 0.1% KOH at 4°C until imaged. Fish were imaged on a Leica M165 FC stereomicroscope. Images were adjusted and white balanced in Adobe Photoshop.

### Kaede photoconversion and imaging

*ptch2:Kaede* fish were anesthetized and placed on walled coverslips as described above. Fins were imaged and viewed on a Nikon Eclipse T*i-*E widefield microscope. A distal portion of the dorsalmost rays was photoconverted using a 405 nm wavelength laser for 1 minute. Immediately after conversion, images were acquired to confirm complete conversion of Kaede from nm to nm emission in the region of interest. Finally, fish were returned to system water and then re-imaged 20 hours later.

### Frozen section immunostaining

Frozen longitudinal sections of 4 dpa fin regenerates were collected as detailed in (Stewart et al., 2014). Sections were placed in Target Retrieval Solution (Dako, S1699) and cooked for 5 minutes in a Cuisinart pressure cooker set to high. Following this, they were blocked for 1 hour at room temperature in 1x PBS, 0.1% Tween-20, 1% DMSO, 1% BSA, 1% NGS block solution. Next, they were left in a solution of primary antibodies diluted in block buffer overnight. The following primary antibodies were used: anti-GFP (1:3000; AVES, GFP-1020), anti-Fbn (1:100; DSHB, JB3), anti-Laminin (1:200; Sigma-Aldrich, L9393) and anti-Runx2 (1:500; SCBT, SC-101145). The next day, sections were treated with a 30 minute high salt wash (0.5M NaCl) followed by washes in PBST. Then, they were stained with Alexa Fluor secondary antibodies diluted 1:1000 in blocking buffer for 1 hour. Finally, they were washed in PBST, nuclear stained using Hoescht, and mounted in SlowFade^TM^ Diamond Antifade (Thermo Fisher S36972). Stained sections were imaged using the Nikon Eclipse T*i-*E with Yokogawa CSU-W1 spinning disk attachments. Images were processed using Fiji and Adobe Photoshop.

### Paraffin section immunostaining

Fins were dissected at 4 or 5 days and prepared as described in (Stewart et al., 2014). Sections were de-waxed in xylenes, then rehydrated. After a quick wash in PBS + 0.1% Tween 20 (PBST), sections were moved to antigen retrieval solution (1mM EDTA pH 8.0 + 0.1% Tween 20) and cooked for 5 minutes in a Cuisinart pressure cooker set to high. Following this, they were blocked for 1 hour at room temperature in 1% milk in PBST. Next, they were left in a solution of primary antibodies diluted in block buffer overnight. The following primary antibodies were used: anti-GFP (1:3000; AVES, GFP-1020), anti-α-Catenin (1:1000; GeneTex,GTX106014), and anti- Runx2 (1:100; LS Bio, LS-B4293-100). The next day, sections were treated with a 30 minute high salt wash (0.5M NaCl) followed by washes in PBST. Then, they were stained with Alexa Fluor secondary antibodies diluted 1:1000 in blocking buffer for 1 hour. Finally, they were washed in PBST, nuclear stained using Hoechst, and mounted in SlowFade^TM^ Diamond Antifade (Thermo Fisher, S36972). Stained sections were imaged using the Nikon Eclipse T*i-*E with Yokogawa CSU- W1 spinning disk attachments. Images were processed using Fiji and Adobe Photoshop.

### In situ hybridization

Probes to detect *fbn3* mRNA were designed and synthesized by ACD Bio. RNAscope^TM^ in situ hybridization was performed using the Multiplex Fluorescent kit (ACD Bio, 323100) according to the manufacturer’s recommendations for paraffin wax embedded sections with minor modifications. *fbn3* probe was diluted 1:100 and the Opal 570 (ACD Bio, PN FP1488001KT) secondary probe was diluted 1:2000. Following RNAScope^TM^, *shha:GFP* sections were antibody stained for GFP with anti-GFP (1:500; AVES, GFP-1020).

### RT-qPCR

Tissue was collected from 3 pools of 3 age matched ABC and *fbn3^b1468/b1468^* fish at 4 days post amputation into TRI Reagent (Zymo Research, R2050-1-200). A VDI 12 Homogenizer (VWR, 82027-184) was used to homogenize tissue, then RNA was extracted and purified using the Direct-zol RNA miniprep (Zymo Research, R2050). cDNA was generated from total RNA using the Maxima^TM^ H-Minus First Strand cDNA synthesis kit (Thermo Fisher, K1651). qPCR was done on cDNA using KAPA^TM^ SYBR FAST qPCR kit (KAPA Biosystems, KK4605) in the StepOne Plus Real Time PCR (Thermo Fisher, 43-766-00). Primer sets spanning introns were used for *fbn1*, *fbn2*, and *fbn3*. Relative expression levels were determined by normalizing to *rpl8.* Primers for *fbn1*, *fbn2*, and *fbn3* were designed in (De Rycke et al., 2025).

## ACKNOWLEDGEMENTS

We thank the University of Oregon AqACS facility for zebrafish maintenance; Dr. Victor Lewis for bioinformatics assistance; the UO zebrafish community and the Stankunas lab for feedback.

## FOOTNOTES

### Author contributions

S. H., G. Y., and K. S. designed experiments with input from S. S., S. H., G. Y., and A. H. performed experiments. S. H. and K. S. prepared and wrote the manuscript with the other authors’ input.

## Data and material availability

Requests for materials should be addressed to K. S.

