## Supplementary Information for "Sonic hedgehog signaling promotes basal epidermal *fibrillin 3* expression for zebrafish fin ray branching"

**Supplemental Materials & Methods**

**Supplemental Figures 1-8**

**Supplemental Materials & Methods**

**Guide RNA protospacer sequences:**

| Target | Exon Number | Protospacer Sequence |
|---|---|---|
| cyp26c1 | Exon 1 | GGGACAGCAAAGCGGACACG |
| cyp26c1 | Exon 2 | GATGCGGAAGGTGAGTGACT |
| crispld2 | Exon 1 | GGAGCTCCAGGGAATCGGGG |
| crispld2 | Exon 2 | GGTGAGGTCCATGTTCCCAT |
| col21a1 | Exon 1 | GGTTGGTACTTCAAGAACAT |
| col21a1 | Exon 2 | GATCGGTCAAAAGTTCACCC |
| fbn2b | Exon 1 | GGCTTGCACCGAATGGACTG |
| fbn2b | Exon 2 | GGTACTTACGAACTATACAC |
| fgfr1bl | Exon 1 | GTGCTGTGTCTACACGAGAA |
| fgfr1bl | Exon 2 | GGCGTGTCCGTCATTCCAAG |
| ncana | Exon 2 | GTCGGCAGTGATGGCGTCGG |
| ncana | Exon 3 | GGCGTCCTGCGAATGCCTGG |
| si:ch211_106n13.3 | Exon 1 | GGCTCCGTCTCATACCTGGG |
| si:ch211_106n13.3 | Exon 2 | GGAACGGTGAACGATCAGAA |
| slit2 | Exon 1 | GGGACAGCGGTGGACTGCCA |
| slit2 | Exon 2 | GGAATACTTACAGAACTCGA |
| slitrk2 | Exon 1 | GGGTATTGGGAATCAGATGG |
| slitrk2 | Exon 1 | GAAGAACCGGTGCTCCTGCG |
| smpd3 | Exon 1 | GATCCAATAGCCAGTAGCAA |
| smpd3 | Exon 3 | GGTCGAAGTTGAGGTCTCCG |
| thsd7ba | Exon 3 | GTTCGTTGCATGCGCAGCGA |
| thsd7ba | Exon 5 | GATGAGGGGTGGAATTGTCT |

**PCR Primers:**

| Target | Forward Primer | Reverse Primer |
|---|---|---|
| *fbn3* Exon 2 | GCTTCCACTCCTATTGTTGTCC | ACCTTGTGTTGTGGTTAAAGGG |
| *fbn2* Exon 2 | TGGCATGTGGAGAATTATTTT G | CACCCAATGAACTGAGAGAGT G |
| *thsd7ba* Exon 3 | CATTGTGTCCGATTTCTCCTCT | TTCTTTCGGTATCTCCTGTGCT |
| *thsd7ba* Exon 5 | GGACAGAAGGCTTGACAATGT T | TTCTACAGTGGAACGAAGTGG A |

### Supplemental Figures

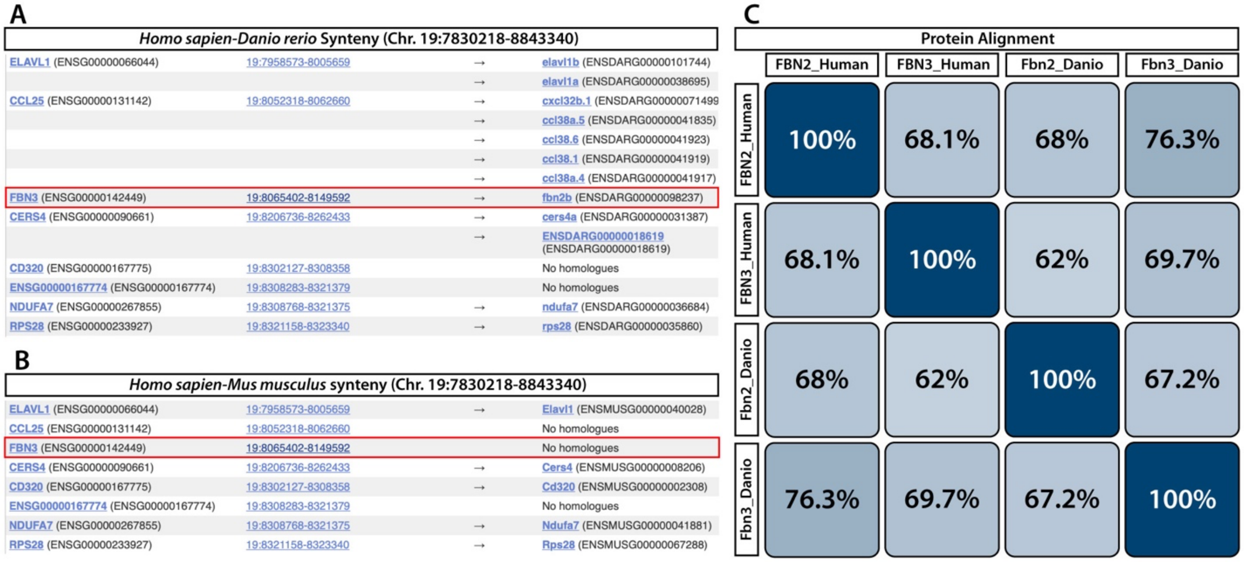


**Supplemental Figure 1.** ***fbn3*** **shares synteny with** ***FBN3*** **and** ***fibrillin 3*** **is absent from the mouse genome. (A)** Synteny comparison between the human and *Danio rerio* genomes showing *FBN3* and neighboring genes. *FBN3* shares synteny with zebrafish *fbn3 (fbn2b)*, as highlighted. **(B)** Synteny comparison between the human and mouse genomes. *FBN3* is absent from the mouse genome. Both analyses were performed using the Ensembl genome browser. (C) Percent sequence homology between human FBN3 and FBN2 and zebrafish Fbn2 and Fbn3 proteins. Analysis was done using Uniprot protein alignment.

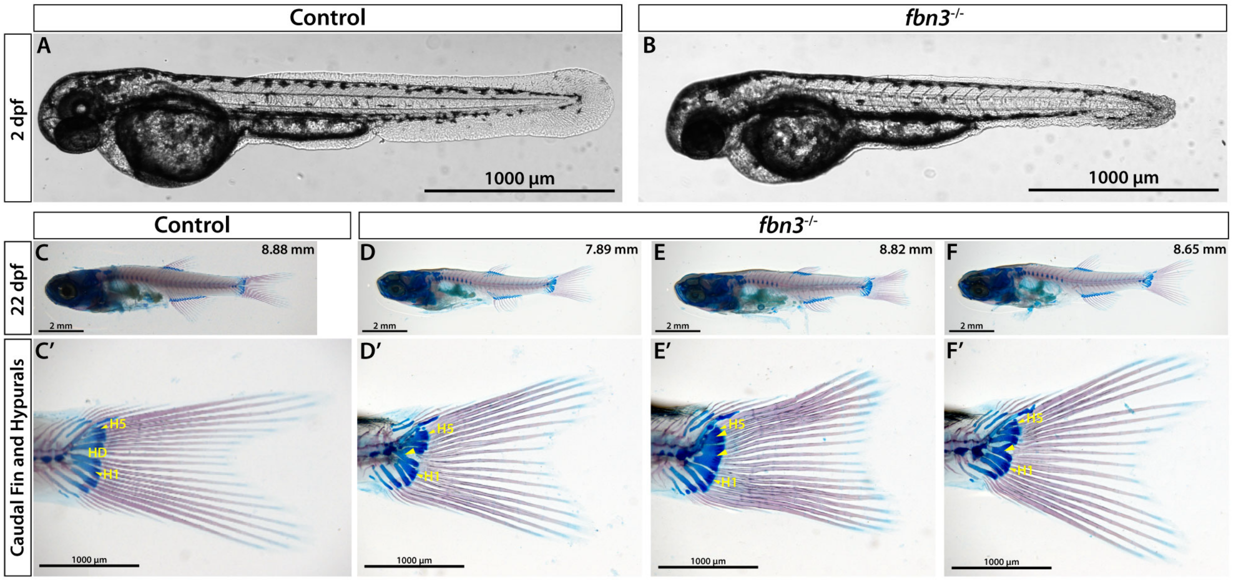


**Supplemental Figure 2. Fibrillin 3 supports fin fold formation and patterning of caudal endochondral structures in zebrafish. (A)** 2 day post fertilization (dpf) control sibling. (B) 2 dpf *fbn3*$^{-/-}$ fish. **(C)** Image of a 22 dpf sibling control stained with Alcian Blue and Alizarin Red (ABAR). Standard length is indicated in the top right corner. (C') Caudal fin image corresponding to the fish in (C). Hypural 1 (H1) and hypural 5 (H5) are labeled in yellow, and the hypural diastema (HD) is denoted at the center of the fin. **(D-F)** *fbn3*$^{-/-}$ fish stained with ABAR. Standard length is indicated in the top right corner of each panel. (D') Caudal fin image of the fish in (D). The yellow arrow indicates a fusion between the bases of hypurals 2 and 3 bridging the hypural diastema. (E') Caudal fin image corresponding to the fish in (E). The yellow arrow indicates a complete fusion of hypurals 2 and 3. (F') Caudal fin image from the fish in (F). The yellow arrow indicates an ectopic extension from hypural 2.

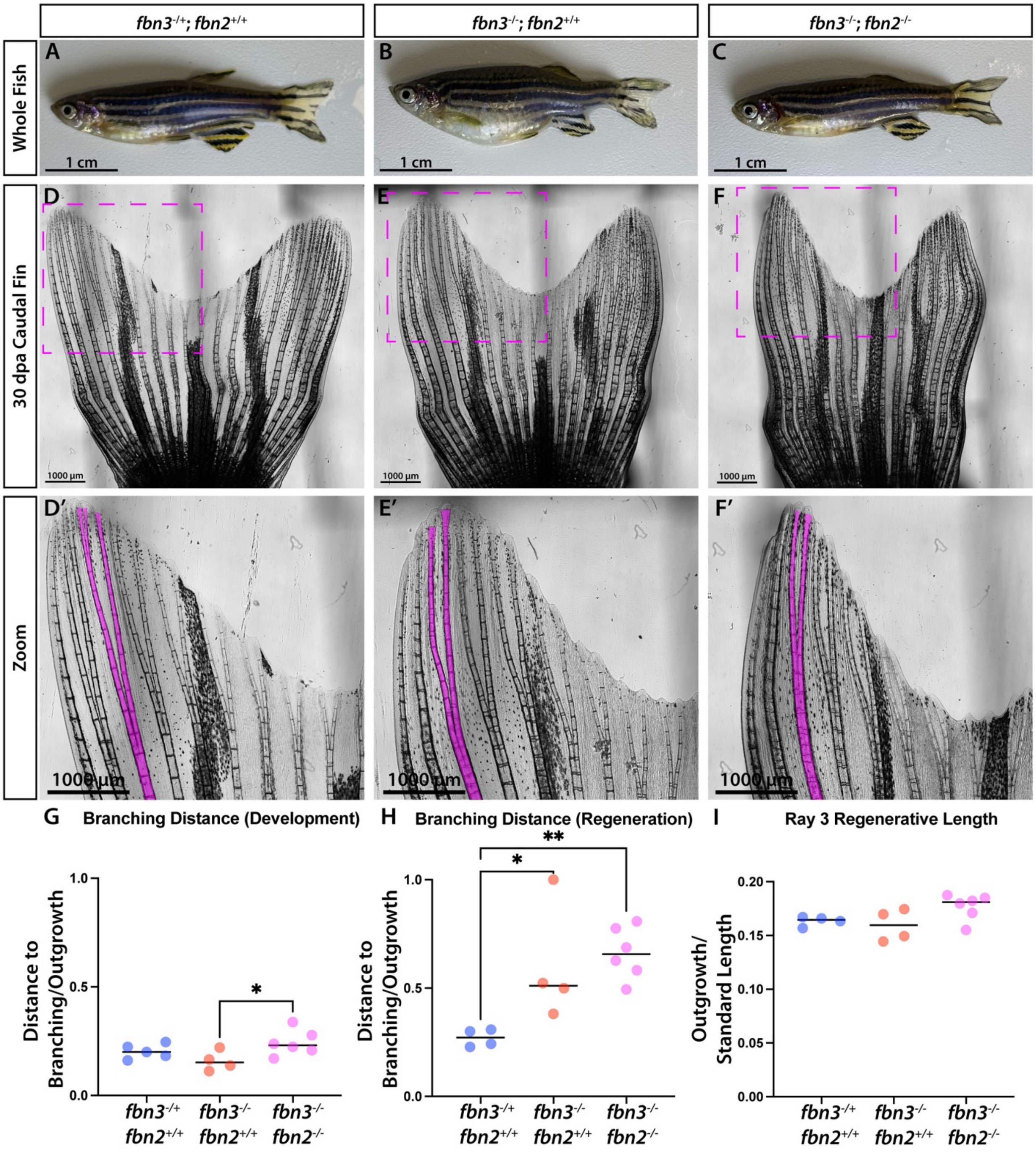
fbn3-/+; fbn2+/+
fbn3-/-; fbn2+/+
fbn3-/-; fbn2-/-
Whole Fish
A
B
C
1 cm
30 dpa Caudal Fin
D
E
F
1000 µm
Zoom
D′
E′
F′
1000 µm
G Branching Distance (Development)
H Branching Distance (Regeneration)
I Ray 3 Regenerative Length
Distance to Branching/Outgrowth
Outgrowth/ Standard Length
1.0
0.5
0.0
0.20
0.15
0.10
0.05
0.00
*
**
fbn3-/+ fbn2+/+
fbn3-/- fbn2+/+
fbn3-/- fbn2-/-

**Supplemental Figure 3. Fibrillin 2 supports Fibrillin 3 to promote ray branching morphogenesis. (A-C)** Images of *fbn3*$^{+/-}$; *fbn2*$^{+/+}$ controls, *fbn3*$^{-/-}$, and *fbn3*$^{-/-}$; *fbn2*$^{-/-}$ fish, respectively. **(D-F)** Regenerated caudal fin images at 30 dpa. Magenta boxes indicate regions shown in zoom panels. (D'-F') Zoomed views of each caudal fin; ray 3 is highlighted in magenta. **(G-H)** Ratio of branchpoint distance to total ray outgrowth for ray 3 during development (G) and regeneration (H). **(I)** Regenerative outgrowth of ray 3 normalized to standard length. Experiments were repeated twice with 5 controls, 4 *fbn3* mutants, and 6 double mutants. Significance determined by one-way ANOVA with Tukey's post-hoc test: $P < 0.05$ (*), **P** < 0.005 (**). Zooms of control and double mutant fins (D', F'), as well as quantifications of branching distance during development and regeneration. (D', F', G, H) are also shown in Figure 6.

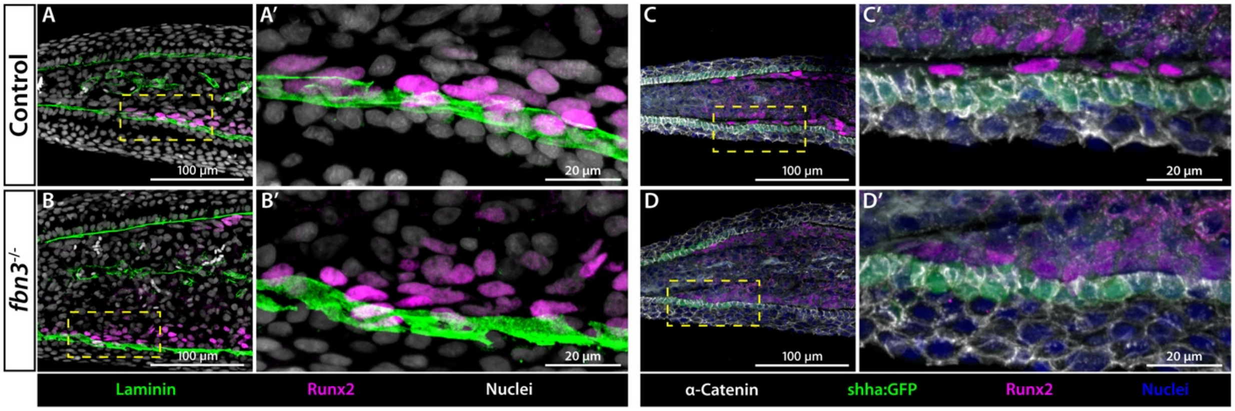


**Supplemental Figure 4. Fibrillin 3 does not regulate basement membrane formation that separates otherwise closely apposed bEp and pOb layers in distal fin regenerates. (A-D')** Immunofluorescent confocal images of longitudinal sections of 4 dpa regenerated caudal fins from (A, C) control and (B, D) *fbn3*$^{-/-}$ fish. (A, B) *Laminin* is in green, Runx2+ pObs in magenta, and nuclei in grayscale. (C, D) α-catenin is in grayscale, *shha:GFP*-expressing bEps are in green, Runx2+ pObs in magenta, and nuclei in blue. The yellow dashed boxes mark the regions shown in corresponding zoom panels. Each experiment was repeated twice, each time using three control and three mutant samples.

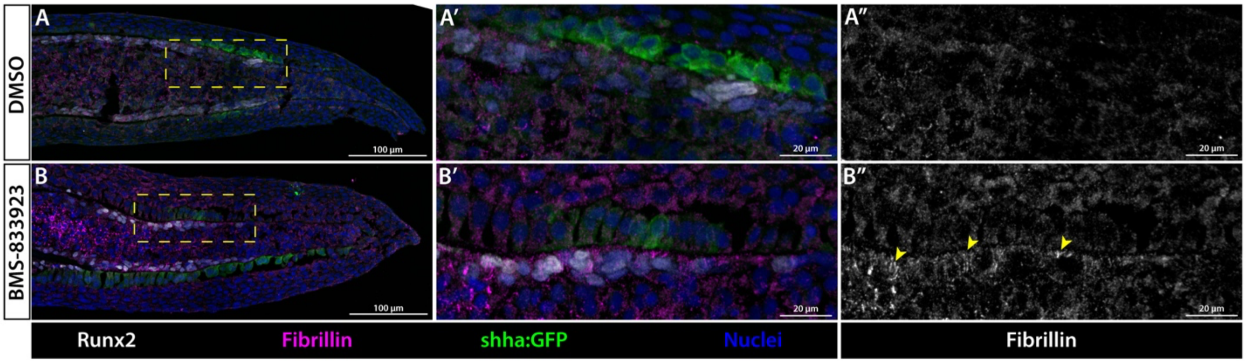


**Supplemental Figure 5. Sonic Hedgehog signaling inhibits the formation of Fibrillin microfibrils at the interface between bEps and pObs in distal fin regenerates. (A–B)** Confocal immunofluorescence images of longitudinal sections from 4 dpa *shha:GFP* caudal fin regenerates. (A) DMSO control and (B) BMS-833923-treated, Smoothened-inhibited fish are shown. Fibrillin is in magenta, *shha:GFP* bEps in green, Runx2+ pObs in white, and nuclei in grayscale. Yellow dashed boxes outline the zoomed regions shown in (A′, B’). The same region is shown in grayscale for Fibrillin alone in (A″, B”). The yellow arrows indicate Fibrillin accumulating at the bEp–pOb interface. The staining was repeated twice, with independent experiments each using three control and three BMS-treated samples.

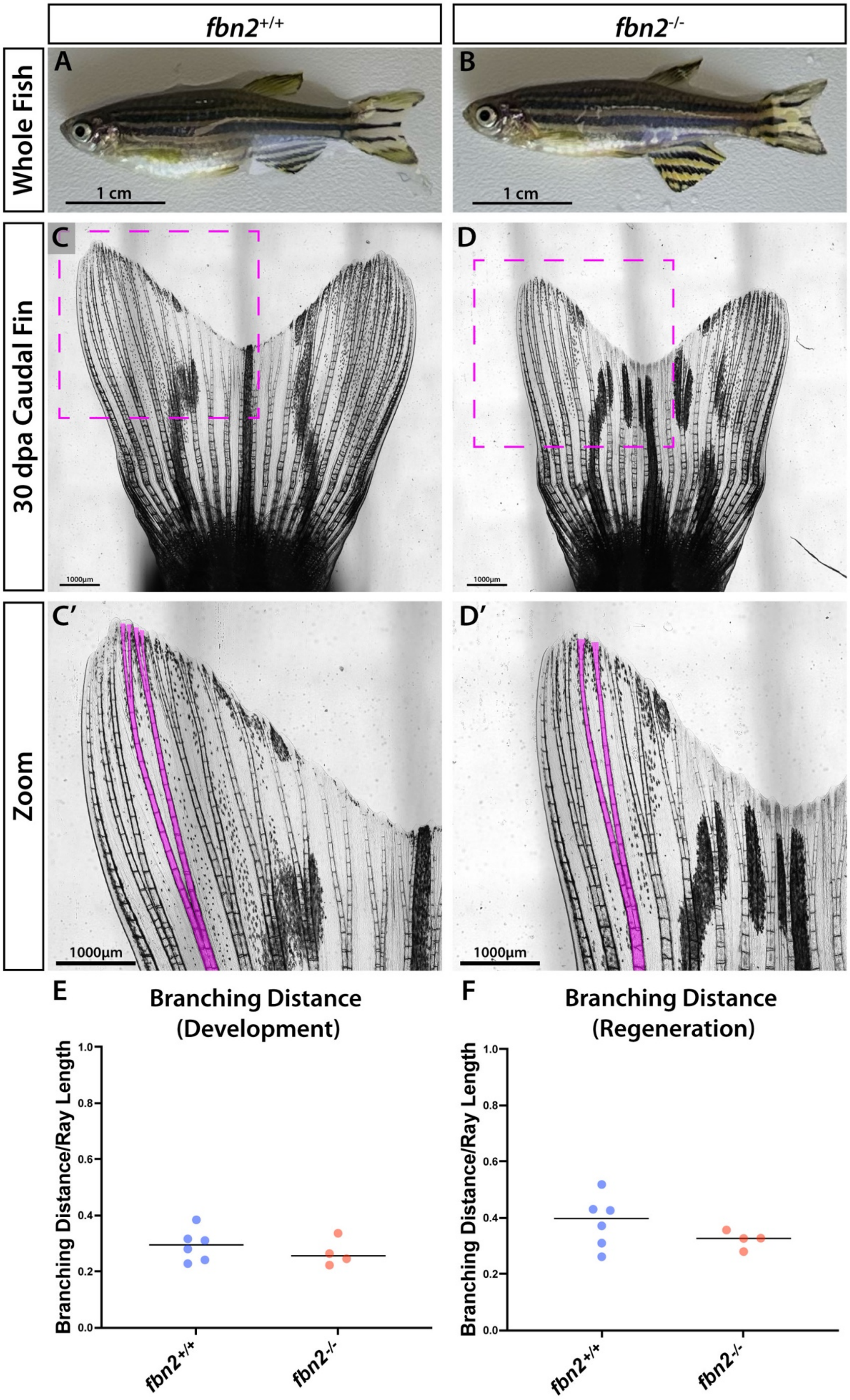

fbn2+/+
fbn2-/-
Whole Fish
A
B
1 cm
1 cm
30 dpa Caudal Fin
C
D
1000µm
1000µm
Zoom
C′
D′
1000µm
1000µm
E
Branching Distance (Development)
F
Branching Distance (Regeneration)
Branching Distance/Ray Length
1.0
0.8
0.6
0.4
0.2
0.0
fbn2+/+
fbn2-/-

**Supplemental Figure 6. *fbn2* is not required for ray branching in regenerating fins.** (A, B) Images of control and $fbn2^{-/-}$ sibling adult fish. (C, D) Caudal fins at 30 dpa. Magenta boxes indicate zoomed distal regions shown in (C′, D′). Ray 3 is highlighted in magenta. (E, F) Ratio of branchpoint distance to total outgrowth in ray 3 during development (E) and regeneration (F). Experiments were repeated twice with six controls and four *fbn2* mutants. Significance was determined by one-tailed unpaired Student's *t*-test: $P < 0.05$ (*).

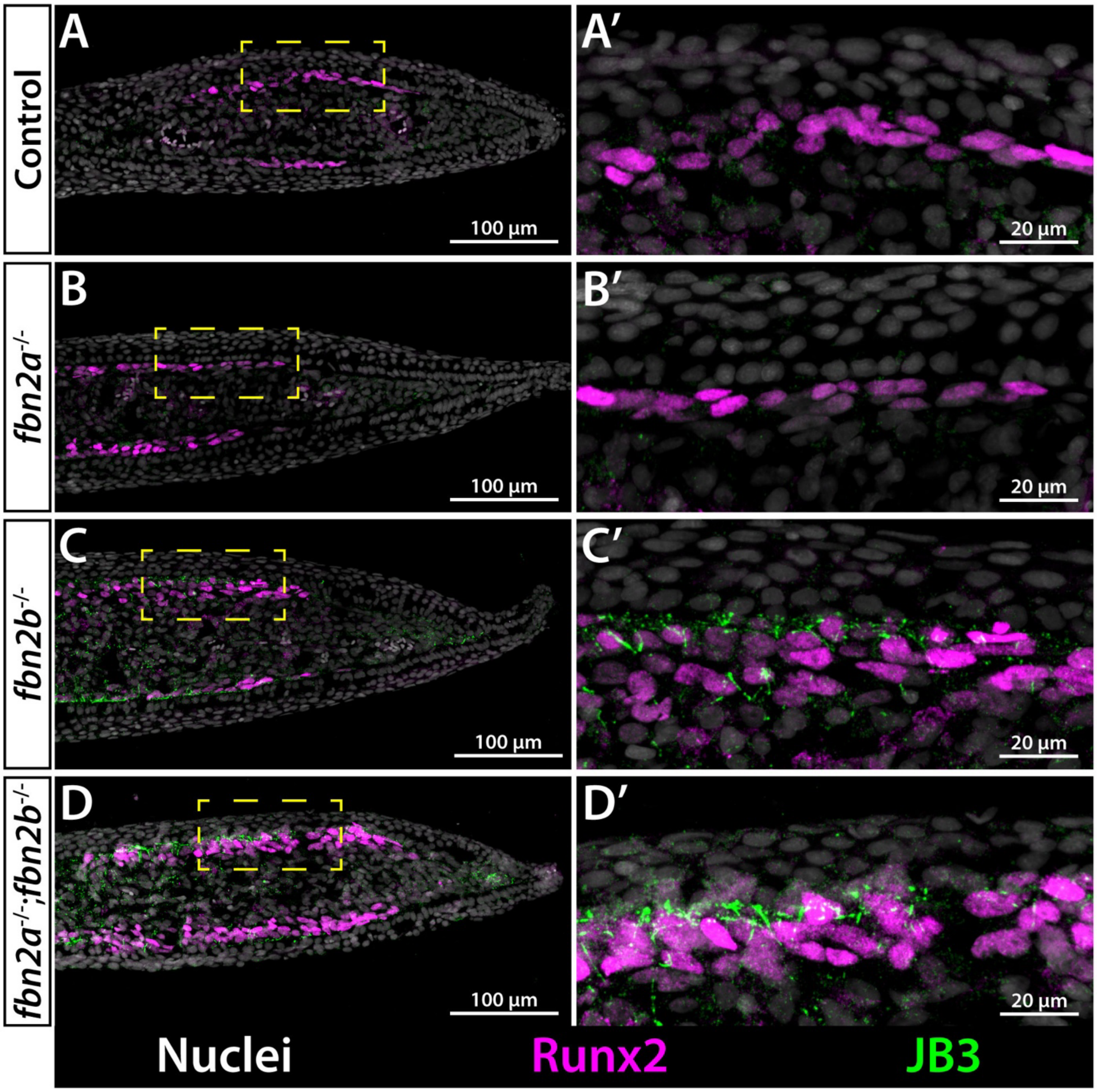


**Supplemental Figure 7. Fbn3 functions with Fbn2 to suppress Fbn1 microfibrils in distal caudal fin regenerates. (A-D')** Confocal, immunofluorescent images of 4dpa longitudinal caudal fin regenerate sections from (A) *fbn*$^{+/-}$;*fbn2*$^{+/+}$ control, (B) *fbn2*$^{-/-}$ , (C) *fbn3*$^{-/-}$, and (D) *fbn3*$^{-/-}$*; fbn2*$^{-/-}$ fish. Fibrillin 1, stained with JB3 antibody, is in green, Runx2-expressing pObs are in magenta, and Hoechst-stained nuclei are in grayscale. Dashed yellow boxes indicate regions shown in the adjacent zoom panels. (A', B', C', D') are also shown in Figure 6. Experiments were repeated twice.

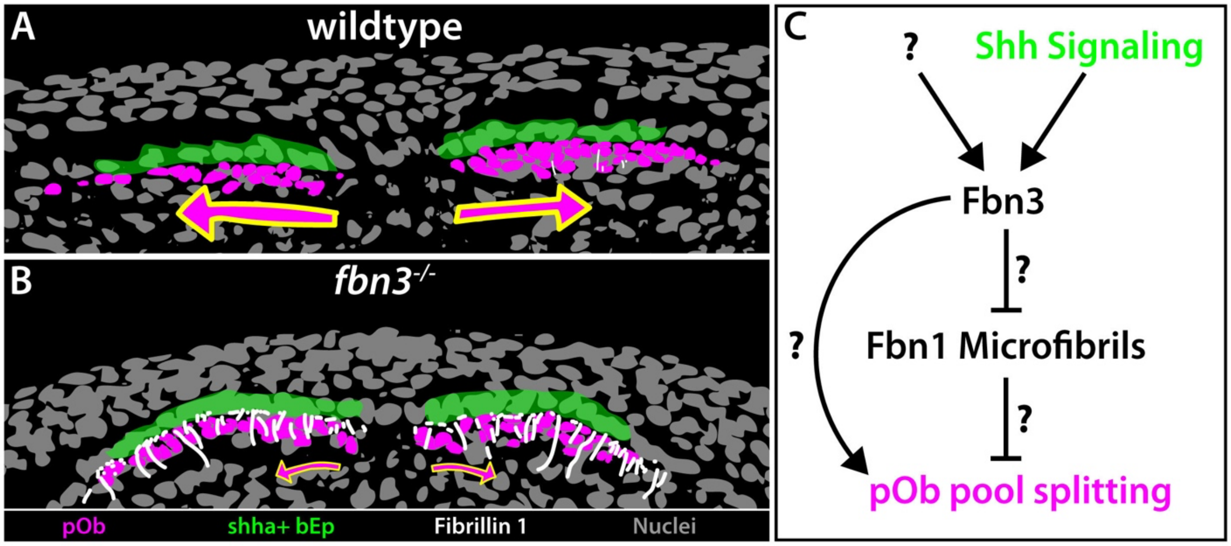


**Supplemental Figure 8. Fbn3 regulates microfibrils in distal fin regenerates to maintain an environment permissive for pOb pool splitting.** (A) Transverse section illustration of a branching ray in a control fish. Magenta arrows indicate lateral pOb movements that gradually split the pOb pools for ray branching. Green highlighting denotes *shha*-expressing bEps and magenta denotes pObs. (B) Transverse section illustration of a branching ray in an *fbn3* mutant. Smaller magenta arrows indicate restricted lateral migration of pObs, possibly due to denser Fibrillin 1 microfibrils (white). (C) Pathway model depicting the role of Shh signaling on Fibrillin 3, Fibrillin 1 microfibrils, and pOb pool splitting. Fbn3 (with Fbn2), including downstream of Shh signaling in bEps, suppresses Fbn1-containing microfibrils, to support pOb movements underlying pOb pool splitting. Locally-produced Fbn3 could also directly support pOb ray branching movements. Shh signaling likely has additional target genes, including in pObs. Nuclei in illustrations were generated by image tracing Hoechst-stained nuclei using Adobe Illustrator. *shha:GFP*-expressing bEps, pObs, and microfibrils were traced from immunofluorescent sections.
